# Anterior default mode brain state dynamics predict depressive symptom severity before and during TMS treatment

**DOI:** 10.64898/2026.05.27.728312

**Authors:** Carina Forster, Chetan Gohil, Bjorn Burgher, Ilya Kuzovkin, Mats W.J. van Es, Mark Woolrich, Diego Vidaurre, Martijn P. van den Heuvel, Cameron Higgins, Luca Cocchi

## Abstract

Transcranial magnetic stimulation (TMS) targeting the left dorsolateral prefrontal cortex is known to progressively reduce symptoms of depression. However, the neural mechanisms supporting this effect are poorly understood. To address this gap, we analysed longitudinal EEG recordings from 70 people undergoing TMS therapy and fitted an established dynamic network model of resting-state activity. Greater baseline symptom severity was associated with reduced occupancy of and fewer transitions into an anterior default mode brain state, alongside increased activity in a posterior default mode state. During treatment, decreases in anterior default mode state engagement following TMS predicted symptom improvement in the latter half of the intervention. Brain state activity exhibited structured, cyclical dynamics, with slower cycles linked to greater baseline severity. These findings suggest that symptoms of depression are characterised by gradual alterations in brain state dynamics, highlighting a central and dissociable role of default mode brain states in the persistence and remission of symptoms.

## Introduction

Transcranial Magnetic Stimulation (TMS) is an effective treatment for refractory depression^1–3^. However, clinical responses to TMS vary substantially across individuals. This variability is thought to reflect differences in patient characteristics^4^, stimulation parameters^5^, and the ability to effectively modulate brain networks supporting symptoms^6,7^.

Neuroimaging studies consistently implicate brain networks, particularly the default mode (DMN) and executive control networks, in the pathophysiology of depressive disorder and treatment response to TMS^7–12^. Evidence obtained by combining neuromodulation and neuroimaging approaches suggests that prefrontal regions encompassing executive and salience brain networks exert regulatory control over the medial prefrontal node of the DMN^13^. Disruptions in these cross-network interactions, involving the medial prefrontal, subgenual cingulate, and insula regions, are a hallmark of depression and are modulated by antidepressant interventions such as TMS^14–16^. Notably, the therapeutic effects of TMS on the left dorsolateral-prefrontal cortex (DLPFC) are thought to be mediated by the correlation between brain activity in this region and the subgenual cortex^2,8,17^. However, existing studies characterise treatment effects using single pre-treatment and post-treatment measurements, typically acquired weeks apart. These designs cannot resolve how brain activity evolves during TMS therapy, nor capture the immediate neural modulation induced by individual TMS sessions^18,19^.

The ability to map changes in large-scale brain dynamics^20,21^ during TMS treatment represents a critical step toward understanding the link between neuromodulation effects on the brain and clinical improvement. Neural markers obtained from scalp electroencephalography (EEG) have been shown to differentiate depression subtypes^22^ and predict treatment response^23^. Initial work has further suggested that temporal changes in EEG-microstates can track depressive symptoms following TMS treatment^24^. Of particular interest is the recent observation that transitions between brain states at rest are not random but are organised into recurring, cyclical patterns^25^. These brain states are characterised by distinct spectral power and coherence profiles that capture coordinated neural activity^20^. The cycles of these brain states reflect fundamental properties of large-scale neural activity and network coordination, features known to be disrupted in depression^26,27^.

The above findings underscore the potential of scalp EEG to capture clinically meaningful changes in brain activity during neuromodulation therapy, thereby enabling a deeper understanding of the neural basis of depression and the optimisation of TMS protocols based on the dynamic expression of cortical activity patterns. Given that TMS of the DLPFC engages fronto-parietal and default mode circuitry^28,29^, we hypothesised that therapeutic effects would be associated with changes in the expression of default-mode and executive brain states. Specifically, we predicted that TMS would modulate the dynamics of these states within and across TMS sessions and that changes in symptom severity could be predicted by changes in state occupancy, transitions, and cycle dynamics across treatment.

To test these hypotheses, we implemented a longitudinal design with six resting-state EEG recordings acquired before and after TMS sessions during fMRI-personalised, robotically delivered TMS therapy for depression^2,30^. By combining repeated pre- and post-TMS session measurements with time-resolved modelling of large-scale brain state dynamics, we characterised how neuromodulation perturbs brain state dynamics and how these effects predict symptom changes.

## Results

We analysed a dataset of 70 individuals diagnosed with treatment-resistant depression who underwent fMRI-personalised and robotically-delivered TMS therapy at the Queensland Neurostimulation Centre (Brisbane, Australia)^2,30^ (**Table 1;** details in Methods). Participants received 20-30 weekday sessions of intermittent theta-burst stimulation (600 pulses per session at 120% of resting motor threshold) of the left DLPFC delivered at rest. Resting-state EEG was recorded immediately before and after the first, 11th, and 20th TMS sessions, enabling assessment of acute stimulation-related changes across multiple treatment stages (**Figure 1**). Depressive symptom severity was assessed weekly throughout treatment using the Hospital Anxiety and Depression Scale – Depression subscale (HADS-D)^31^. Sample characteristics are presented in Table 1 and supplementary text.

**Figure 1:**
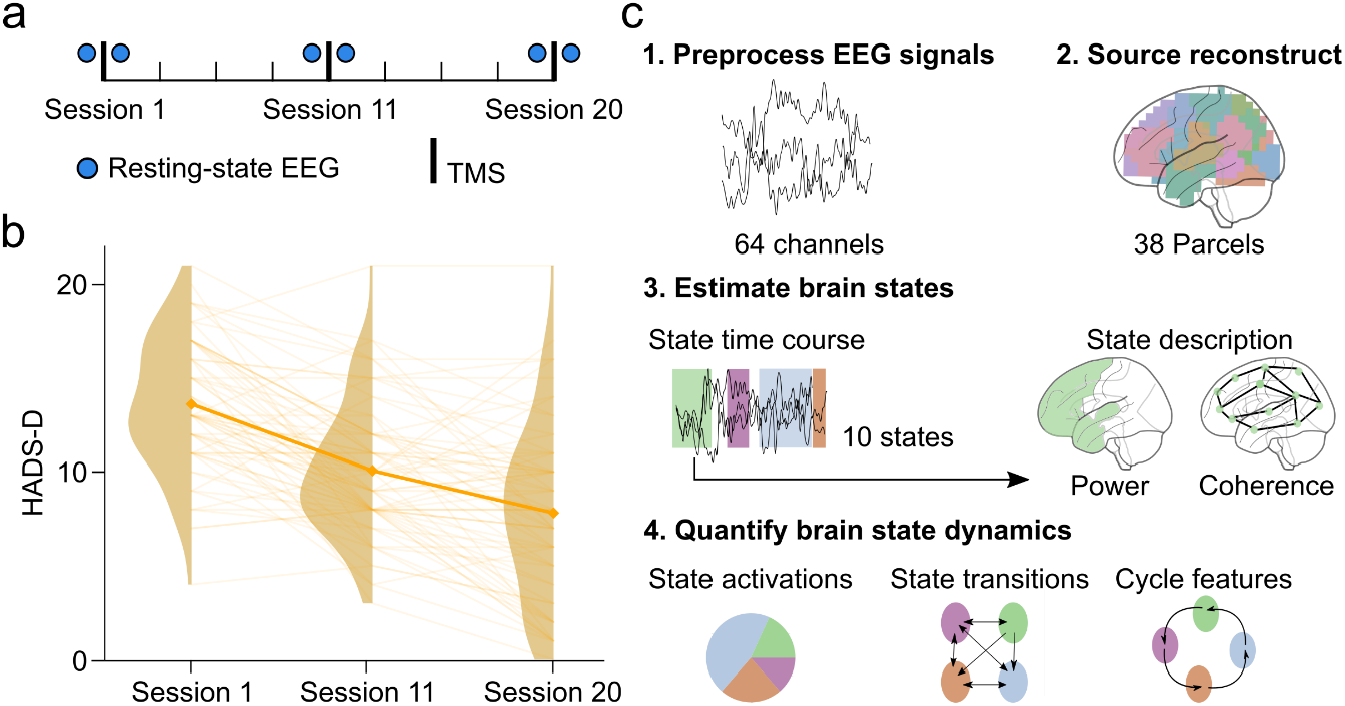
Study timeline, treatment response, and analysis pipeline of resting state EEG. **a**. Participants with refractory depression received fMRI-personalised and robotically delivered repetitive TMS (intermittent theta burst stimulation, iTBS^32^) to the left DLPFC over the course of 4 weeks (1 session a day for 4 weeks, 5 days a week). Resting-state eyes-closed EEG was collected at Session 1, Session 11, and Session 20, with one recording immediately before and one recording immediately after each TMS session (denoted with blue dots). **b**. Self-reported depressive symptoms (HADS-D^31^) significantly improved from Session 1 to Session 20 (p < 0.001) (see main text, details in Supplementary Table 3). **c**. Resting-state EEG analysis pipeline: 1. Raw eyes-closed resting-state EEG data from each session were preprocessed using a semi-automated pipeline implemented in the osl-ephys^33,34^ and MNE-Python^35^ packages. 2. The clean EEG data were source localised using an individual MRI scan and 38 cortical parcels. 3. 10 brain states were estimated using a canonical time-delay embedded Hidden Markov Model^20,36^. We applied the pretrained model to our EEG data and obtained a state time course for each individual and each session. Each brain state is characterised by a unique cortical topology and spectral profile (power and coherence). 4: State dynamics were quantified, focusing on state activations (how active a state was in each recording) and state transitions (the probability of switching from one state to another). Long-range dynamics^25^ were characterised using cycle features.

**Table 1.** Sample characteristics.

| Measure | Total |
| --- | --- |
| Sample size | 70 |
| Age, years |  |
| Median (IQR) [min-max] | 38.5 (29.0–50.0) [18–77] |
| Gender |  |
| Female | 40 (57.1%) |
| Male | 30 (42.9%) |
| Handedness |  |
| Right | 64 (91.4%) |
| Left | 6 (8.6%) |
| Years of education |  |
| Median (IQR) [min-max] | 14.5 (12.0–17.8) [6–22] |
| Age of MDD diagnosis, years |  |
| Median (IQR) [min-max] | 21.0 (17.0–34.8) [12–57] |
| Age of depressive symptoms onset, years |  |
| Median (IQR) [min-max] | 18.5 (16.0–34.0) [8–57] |
| Research tiers* (1/2/3) | 15(21.4%) / 45(64.3%) / 10(14.3%) |
| Previous rTMS |  |
| Yes | 11 (15.7%) |
| No | 59 (84.3%) |
| Previous ECT |  |
| Yes | 8 (11.4%) |
| No | 62 (88.6%) |
| Treatment regime |  |
| rTMS sessions, n |  |
| Median (IQR) [min-max] | 26.0 (20.0–30.0) [17–30] |
| Days from pre-time point to post-time point |  |
| Median (IQR) [min-max] | 63.0 (53.0–70.0) [37–129] |

\*1: Primary MDD, 2: MDD with complex comorbidity, 3: Bipolar/structural brain disease.
For details, see Hearne et al. (2025)<sup>2</sup>.

Depressive symptoms decreased significantly over sessions (**Figure 1b**). Compared to baseline, scores were significantly lower at week 2 (β = −2.89, SE = 0.41, 95% CI [−3.70, −2.07], p < .001) and week 4 (β = −4.63, SE = 0.41, 95% CI [−5.44, −3.82], p < .001, suppl. fig. 1a shows symptom trajectories from pre to post treatment).

To define brain state dynamics across the treatment, we applied a 10-state canonical^36^ time-delay embedded Hidden Markov Model (TDE-HMM)^20^ (see Methods). Among the 10 brain states characterising resting dynamics, six states support sensory functions (visual, auditory and somatosensory processing), two states support executive processes, and two states are linked to activity in default mode regions^36,37^. Of interest for our hypotheses, State 1 corresponded to the posterior default mode state, which is characterised by increased low-frequency power in occipito-parietal regions and increased coherence between these regions and the frontal cortex relative to the other brain states (**Figure 2c**). Conversely, the anterior default mode state shows increased low-frequency power in anterior frontal regions and low coherence in parieto-temporal regions (**Figure 2d**). States 9 and 10 exhibit lower power in posterior brain regions than the other states and are commonly referred to as executive-or attention-related brain states. Power and coherence profiles of all 10 brain states are visualised in supplementary figure 2 and model summary statistics in supplementary figure 3.

**Figure 2.**
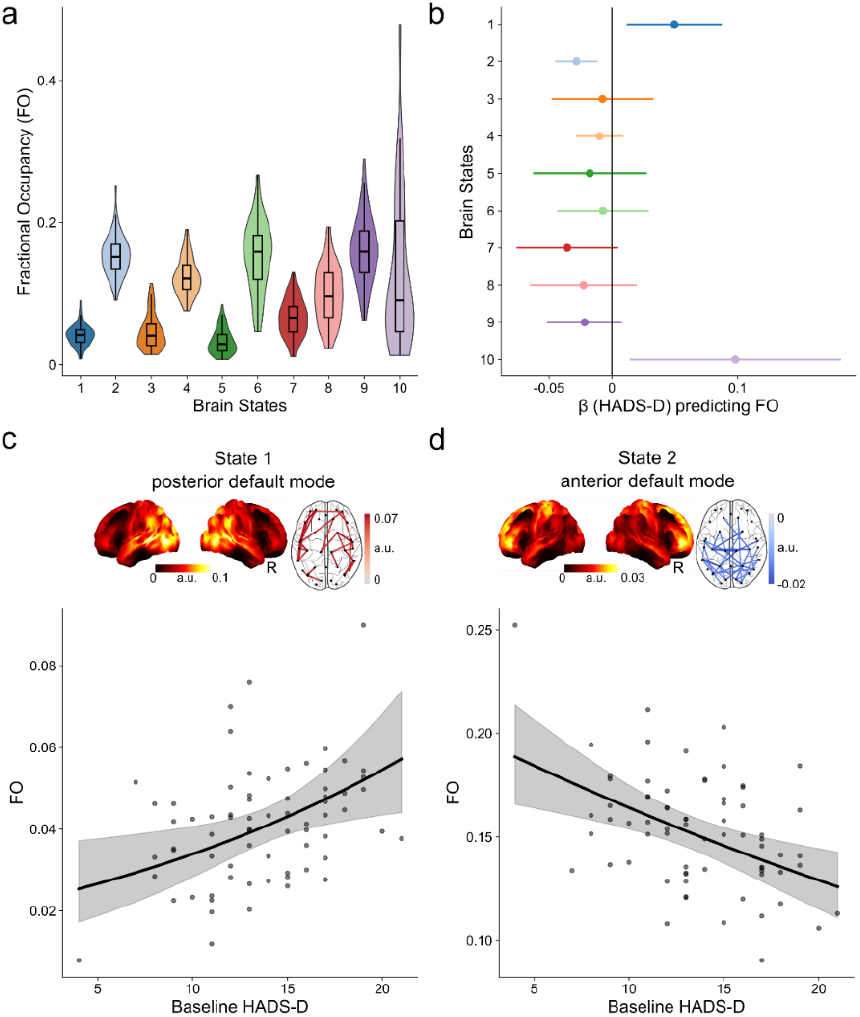
Baseline depression severity is linked to the relative time spent in default mode states. **a**. Fractional occupancy (FO) of each of the 10 canonical brain states at baseline (first EEG recording, before any TMS). FO represents the mean activation of each brain state compared to all other states during the EEG recording. **b**. Results from a multivariate linear regression analysis on fractional occupancy (FO) of each brain state show a significant contribution of baseline depression score (HADS-D) on the expression of brain state 1 (posterior default mode state, p_FDR_=0.043) and brain state 2 (anterior default mode state, p_FDR_=0.008) after controlling for age and gender. **c**. Functional topology and connectivity fingerprint of the posterior default mode network (brain state 1). This state is characterised by relatively higher power in the occipital and parietal cortices and enhanced connectivity between frontal and posterior cortical areas (top 5% strongest connections shown, absolute values). Participants with greater activity in the posterior default mode state show higher baseline depression severity. **d**. Functional topology and connectivity fingerprint of the anterior default mode network. This state is characterised by higher broadband power in the prefrontal cortex and by decreased connectivity between the occipito-parietal cortices. Individuals with a higher baseline depression score showed reduced activation of the anterior default mode state. Points represent observed values; the regression line indicates the model-predicted fractional occupancy, adjusted for covariates. Note that the regression was calculated on logit-transformed FO and back-transformed to original FO values. The shaded area represents the 95% confidence interval of the model predictions. Abbreviations: FO: Fractional Occupancy, HADS-D: Hospital Anxiety and Depression Scale – Depression^31^.

### Baseline depression severity distinctly predicts activation of anterior and posterior default mode brain states

Baseline depression severity measured before TMS treatment predicted fractional occupancy (FO) (i.e., the proportion of time each state is active; **Figure 2a**) in state 1 and state 2 (**Figure 2b**). Individuals with higher symptom severity at baseline showed greater activation of the posterior default mode state (state 1, β = 0.05, 95% CI [0.012, 0.086], SE = 0.019, *p*_*corrected*_ = 0.043, R^2^ = 0.20, N=70 **Figure 2c**) and reduced activation of the anterior default mode state (state 2, β = −0.028, 95% CI [−0.044, −0.012], SE = 0.008, *p*_*corrected*_ = 0.006, R^2^ = 0.22, N=70, **Figure 2d**).

### Acute TMS changes in mid-treatment anterior default mode fractional occupancy are linked to clinical response

We next examined whether acute TMS effects on default-mode network dynamics could predict clinical response to continued TMS therapy. Specifically, we tested whether session-specific pre-to post-TMS modulations in default mode states FO predicted subsequent changes in depressive symptoms following continued TMS therapy.

For the baseline-to-mid-treatment transition, changes in default-mode brain-state FO following TMS (Session 1 pre vs post) did not significantly predict symptoms at Session 11. In the adopted model (R^2^ = 0.38), baseline symptom severity was a significant predictor of symptom severity at mid-treatment (β = 0.579, SE = 0.125, 95% CI [0.334, 0.824], p_uncorrected_ < 0.001), whereas post-TMS changes in FO in both the posterior default mode state (β = −0.067, SE = 0.294, 95 % CI [-0.642, 0.509], p_uncorrected_ = 0.821) and the anterior default mode state (β = -0.313, SE = 0.179, 95 % CI [-0.664, 0.039], p_uncorrected_ = 0.081) were not statistically significant.

Changes in FO linked to the mid-treatment time point (Session 11) significantly predicted symptoms at the final EEG session (Session 20; R^2^ = 0.65). Greater reductions in anterior default mode state occupancy following TMS were associated with lower symptom severity in Session 20 (β = −0.376, SE = 0.146, 95% CI [−0.661, −0.091], p_uncorrected_ = 0.010, **Figure 3**). However, changes in FO in the posterior default mode state were not a significant predictor of symptom improvement (β = −0.296, SE = 0.323, 95 % CI [-0.928, 0.336], p_uncorrected_ = 0.358). Symptom severity at mid-treatment remained a strong predictor of subsequent symptoms (β = 0.770, SE = 0.107, 95 % CI [-0.561, 0.979], p_uncorrected_ < 0.001).

**Figure 3.**
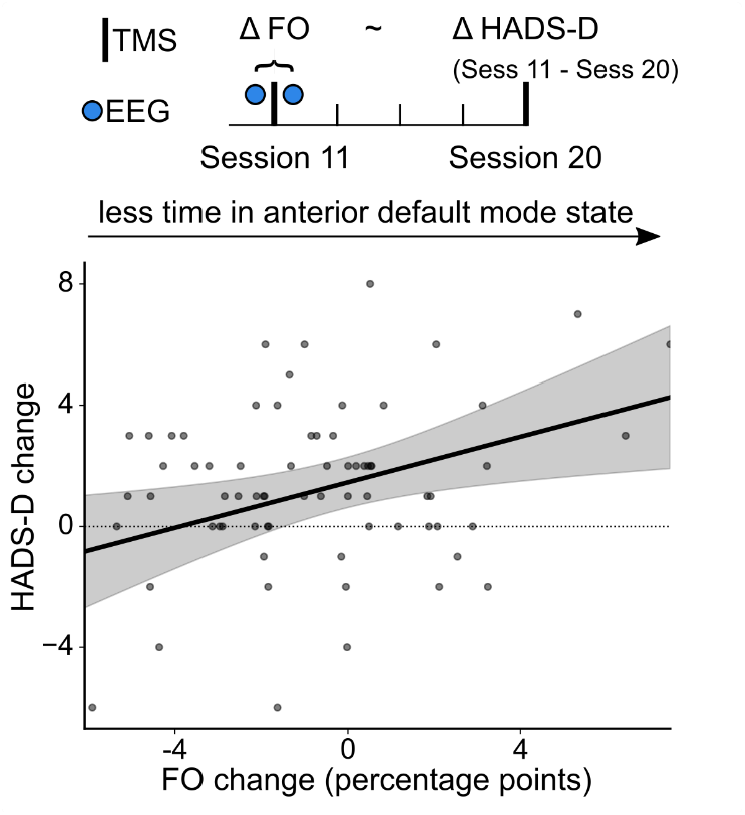
Acute changes in anterior default mode state occupancy following TMS therapy in Session 11 are associated with self-reported depressive symptoms in Session 20. A linear regression model was fitted to data from 70 individuals who completed at least 4 weeks of TMS treatment. The regression model predicted depression severity at Session 20 based on the change in fractional occupancy (FO) of the anterior default mode state in Session 11. Symptom severity at Session 11, age and gender were included as covariates. Reduced time spent in the anterior default mode state at Session 11 following TMS was associated with greater symptom improvement (HADS-D^31^) at Session 20 (p_uncorrected_ = 0.010). FO is expressed as a percentage point change. Points represent observed values; the regression line indicates the model-predicted change in symptoms, adjusted for covariates. The shaded area represents the 95% confidence interval of the model predictions.

### Increased likelihood of transitioning into anterior default mode and attention states is linked to lower baseline symptom severity

We then asked whether the observed changes in state occupancy (FO) were accompanied by altered transitions between states. Examining state transitions offers a more complete characterisation of cortical dynamics during TMS therapy, because individuals may spend comparable amounts of time in the same states yet differ in how frequently they switch between them. We first analysed short-term Markovian dynamics between brain states, modelled by a 10 × 10 Markov state transition probability matrix for each individual and each session (**Figure 4a**). Given the high dimensionality of these matrices, we applied principal component analysis on the transition probability matrices of all sessions and all participants to obtain a low-dimensional representation of state transitions (suppl. fig. 4).

**Figure 4.**
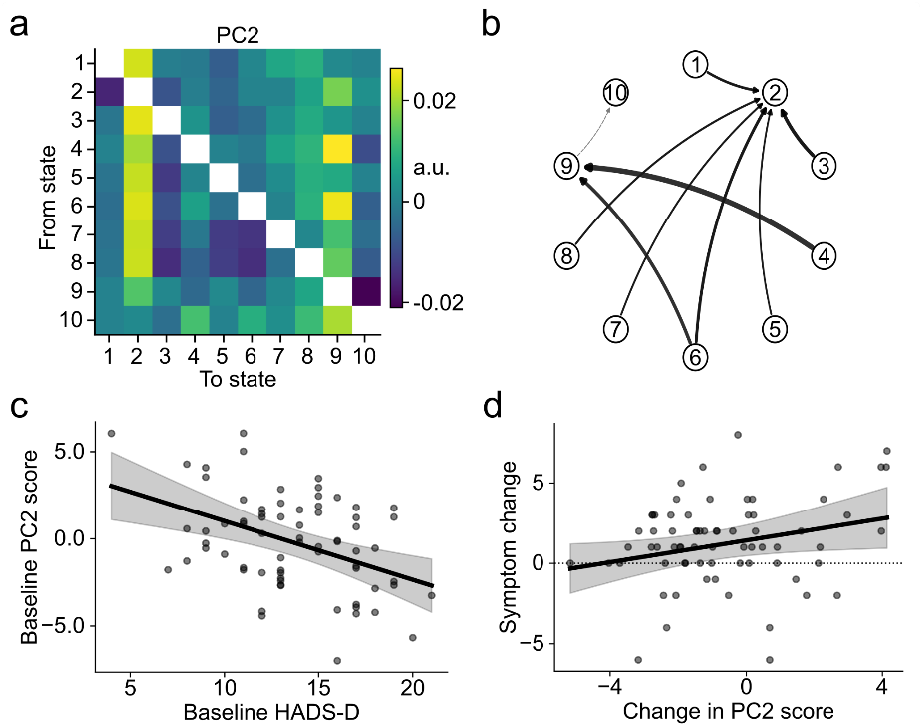
Baseline symptom severity is related to the probability of transitioning into the anterior default mode state and an attention state. **a**. The second principal component (PC2) has strong positive loadings on transitions into the anterior default mode state (state 2) and transitions into an attention-related state (state 9). **b**. Graphical visualisation of the 20 % absolute strongest connections. The thickness of the arrows represents the strength of the transition probability. **c**. Higher PC2 scores obtained from the EEG recording in Session 1 are associated with lower baseline depressive symptoms (p_uncorrected_ < 0.001). **d**. A higher PC2 score pre TMS compared to post TMS (a more positive change in PC2) in Session 11 is associated with a greater change in self-reported depressive symptoms (HADS-D) from Session 11 to Session 20 (p_uncorrected_ < 0.035). Abbreviations: HADS-D: Hospital Anxiety and Depression Scale, Depression subscale^31^.

PC2 was characterised by strong positive loadings on transitions to the anterior default mode state (state 2) and transitions to the attention-related state (state 9, **Figure 4a&b**). PC2 scores were significantly associated with depressive symptom severity, with higher HADS-D scores pre-treatment predicting lower PC2 scores in the first EEG session (Session 1; β = −0.337, SE = 0.093, 95% CI [−0.519, −0.154], p_uncorrected_ < 0.001; R^2^ = 0.193, **Figure 4c**).

PC1 showed strong positive loadings on attention-related states (state 9 and 10) and negative loadings on transitions from the anterior default mode state (state 2, suppl. fig. 5a&b). Depressive symptom severity at baseline was not associated with PC1 component scores in the first EEG session (β = 0.323, SE = 0.237, 95% CI [−0.142, 0.788], p_uncorrected_ = 0.173; R^2^ = 0.07, suppl. fig. 5).

### Acute changes in anterior default mode and attention brain states transitions predict symptom improvement from mid-treatment to the final TMS session

We next examined whether acute TMS effects on default-mode network state transitions could predict clinical response following TMS treatment. Specifically, we tested whether session-specific pre-to post-TMS modulations in PC2 predicted subsequent changes in depressive symptoms.

Baseline to mid treatment symptom improvement did not significantly relate to changes in PC2 scores (β = −0.040, SE = 0.161, 95% CI [-0.356, 0.275], p_uncorrected_ = 0.803, R^2^ = 0.34, suppl. fig. 6). However, clinical improvements between the mid and final EEG session were linked to changes in the transition probabilities captured by PC2. Specifically, greater pre-to post-TMS reductions in PC2 scores at Session 11 were associated with lower symptom severity at Session 20 (β = −0.340, SE = 0.161, 95% CI [−0.654, −0.025], p_uncorrected_ = 0.035; R^2^ = 0.63, **Figure 4d**).

Baseline symptom severity was a strong predictor of symptom change from Session 1 to Session 11 (β = 0.632, p < 0.001) and Session 11 to Session 20 (β = 0.829, p < 0.001).

### Cyclical brain state dynamics are present in depression and linked to baseline symptom severity

Brain dynamics contain both short- and long-term temporal dependencies. Recent work has demonstrated that longer-term brain state dynamics (250 to 1000 ms) in healthy individuals form structured cyclical trajectories at rest, with cycle metrics linked to age, cognition, and behaviour^25^. We therefore asked whether such longer-term cyclical dynamics are present in individuals with clinical depression and whether individual differences in cycle metrics relate to individual symptom severity at baseline and during TMS treatment.

We applied the TINDA algorithm as described by Van Es, Higgins et al.^25^. Results showed that asymmetries in fractional occupancy differ significantly across several brain states (**Figure 5a**), giving rise to a structured pattern of cyclical dynamics across states (**Figure 5b**). This is indicative of the non-random temporal organisation of state transitions. Having confirmed the presence of structured cyclical dynamics in our clinical sample (suppl. fig. 7), we then analysed the same two summary measures previously used to analyse cyclical dynamics in healthy populations: cycle strength and cycle rate^25^. Using regression models, we tested whether baseline depressive symptom severity predicted individual differences in cycle metrics. Baseline depressive symptom severity scores did not significantly predict cycle strength (β = 0.085, 95% CI [−0.128, 0.298], SE = 0.109, p = 0.435; R^2^ = 0.02, **Figure 5c**). However, baseline clinical scores significantly predicted cycle rate (β = −0.045, 95% CI [−0.069, −0.020], SE = 0.012, p < 0.001; R^2^ = 0.21, **Figure 5d**), such that higher baseline depressive symptom severity was associated with a slower cycle rate in the first EEG session.

**Figure 5.**
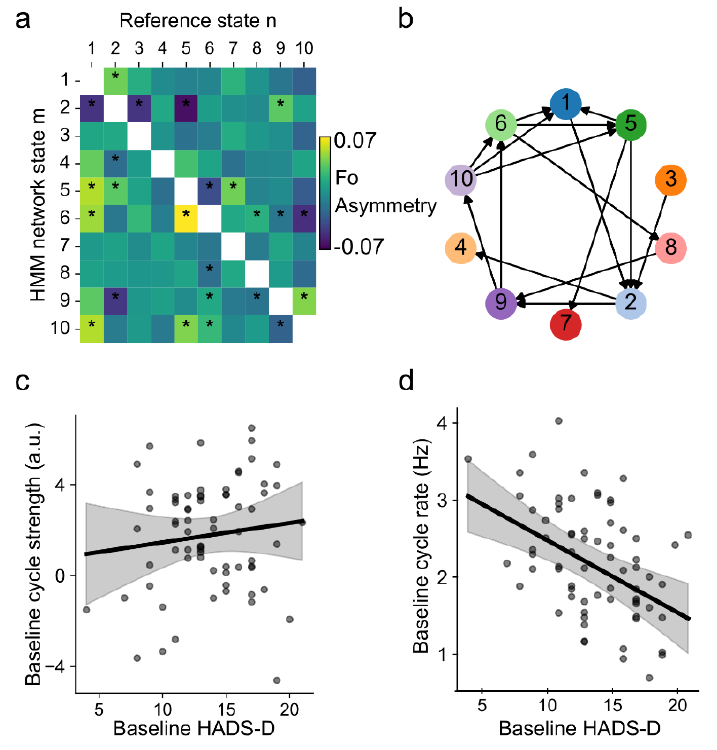
Higher depressive symptom severity predicts lower cycle rate at baseline. **a**. Fractional asymmetry matrix (arbitrary units, a.u.) quantifying directional asymmetries in state transitions; asterisks indicate significant asymmetries (Bonferroni-corrected *p* < 0.05). **b**. Cyclical dynamics^25^ obtained from a ten-state canonical model. **c**. Baseline depression severity was not significantly associated with baseline cycle strength (a.u., *p* = 0.435). **d**. Baseline depression severity was significantly associated with cycle rate (Hz, *p* < 0.001). Individuals with a higher depression severity at baseline had a lower cycle rate at baseline.

We next examined whether acute within-session changes in cycle rate during TMS treatment were associated with subsequent symptom improvement. Changes in cycle rate from pre-to post-TMS did not significantly predict symptom improvement from baseline to mid-treatment (β = −0.210, 95% CI [−1.302, 0.883], SE = 0.557, p = 0.707; R^2^ = 0.35) nor did changes in cycle strength (β = 0.069, 95% CI [−0.114, 0.252], SE = 0.093, p = 0.462). Predicting subsequent symptom improvement from mid-to the final EEG session showed no significant effect of cycle strength (β = 0.011, 95% CI [−0.232, 0.254], SE = 0.124, p = 0.929; R^2^ = 0.61) nor cycle rate (β = −0.360, 95% CI [−0.972, 0.253], SE = 0.312, p =0.250).

#### Control analyses

To assess the dependence of our findings on the adopted brain states resolution (10 states), we repeated all analyses using HMMs with 6 and 8 brain states (suppl. fig. 8, 9, 10).

## Discussion

The aim of this study was to examine whether changes in the dynamic patterns of brain activity during TMS therapy for depression are associated with symptom change. We hypothesised that symptom improvement would be linked to distinct alterations in executive and default-mode brain states. Using longitudinal resting-state EEG recordings and a TDE-HMM^20,21,36^, our study extends findings from prior work by demonstrating that the long-term temporal organisation of brain states contains clinically relevant information. Our analyses partially support the hypothesis that attention-related states underpin symptom severity and improvement during TMS therapy but indicate that activity in default-mode states plays a central role in both, albeit in opposing directions. These findings suggest an important distinct contribution of default mode sub-systems and functions to the persistence of depressive symptoms and the process of recovery.

At baseline, individuals with greater symptom severity spent more time in the posterior default mode state, less time in the anterior default mode state, and had a reduced likelihood of transitioning to the anterior default mode state and an executive brain state. This pattern of brain activity suggests that severe depression is not only characterised by a distinct expression of default mode subsystems^38,39^ but also by reduced ability to access processes supported by the anterior default mode. Activity of anterior default mode regions has been associated with self-referential processing and rumination^10,11,40^ while posterior regions have been shown to support contextual and autobiographical processing^41–43^. Accordingly, our findings support a broad disruption in the balance of internally directed cognition in depression^39^.

While greater baseline symptom severity is associated with reduced anterior default mode engagement, reductions in this state occupancy during TMS treatment predicted greater symptom improvement in the later treatment phase. This effect was evident in both the time spent in the anterior default mode state and the probability of transitioning into this state. These results suggest a functional dissociation within the anterior default mode system, in which its role in the expression of depressive symptoms differs between a relatively stable disease state and a dynamically evolving recovery phase^44^. Reduced baseline engagement may reflect rigid self-referential processes in more severe depression, with the capacity to reconfigure the expression of this state during treatment linked to adaptive neurophysiological^14,29^ or neurocognitive^45^ processes supporting recovery. For example, anterior default mode dynamics during recovery may index the reorganisation of internally directed processing, shifting from maladaptive negative self-referential content towards more adaptive representations^46^.

Although an association between changes in anterior default mode dynamics and subsequent symptom improvement was observed in the first session, it did not reach statistical significance. Because the first EEG session coincides with the initial exposure to TMS pre-to post-changes in the anterior default mode state may be more variable and influenced by nonspecific factors such as high arousal. By the second EEG session (Session 11), nonspecific influences are likely reduced, resulting in more stable and reliable neural and clinical effects. This temporal progression may explain why significant brain-behaviour associations emerge only at later stages of treatment. Early clinical improvements could also reflect nonspecific factors such as expectancy or placebo-related effects^47– 49^ or rapid shifts in cognitive-affective processing that are not yet strongly reflected in large-scale brain dynamics. These findings may explain why reliable neurobiological predictors of TMS response have been difficult to identify^4^, as averaging across treatment stages likely obscures stage-specific effects.

In addition to short-term state dynamics, we observed a structured cyclical organisation of brain state activity, consistent with recent findings in healthy populations^25^. Cycle rate was negatively associated with baseline symptom severity, with slower dynamics observed in individuals with more severe depression. This finding aligns with evidence linking slower brain dynamics to reduced cognitive flexibility^25,50^ and accelerated brain ageing^51^ in depression. However, changes in cyclical dynamics during treatment were not robust predictors of symptom improvement, suggesting that these slower-timescale processes may primarily reflect trait-like aspects of the disorder rather than mechanisms that directly facilitate symptom recovery.

Interpreting our results requires consideration of the heterogeneity of pharmacological treatments received by participants alongside TMS. This may confound the contribution of neuromodulation to brain state activity and symptom improvement. The naturalistic clinical design precluded inclusion of a sham control condition, limiting causal inference on specific TMS effects. Replication in sham-controlled trials will be essential to determine whether the observed neural changes reflect TMS-specific mechanisms.

Overall, this study provides initial evidence that scalp EEG-derived brain-state dynamics serve as informative markers of depressive symptom severity and symptom trajectories during TMS treatment. Specifically, we show that anterior and posterior default mode states capture variation in baseline symptom burden and provide evidence of state-dependent changes in anterior default mode dynamics. These findings suggest that the temporal organisation of default-mode brain states represents a key neurophysiological marker of depressive pathophysiology and a promising target for guiding and optimising neuromodulation-based interventions, including strategies to sustain therapeutic gains following TMS.

## Methods

### Participants

Eighty-five individuals with refractory depression consented to the research procedures while undergoing TMS therapy at the Queensland Neurostimulation Centre (QNC). Neuroimaging-personalised and robotically delivered TMS treatment was authorised by the Australian Therapeutic Goods Administration (TGA: DAP21-0020539). The research protocol was approved by local ethics committees (UnitingCare and QIMRB). All participants provided written informed consent.

Participants were excluded if they had missing clinical data (N = 3) or MRI scans of poor quality (e.g., inadequate field of view) (N = 7). Five additional participants were excluded due to excessive noise in at least one EEG recording. The final research sample comprised 70 participants. Of these, 40 identified as women. The median age was 39 years (range: 18–77 years), and the median duration of depressive illness was 11 years (range: 1–65 years). The number of TMS treatment days ranged from 17 to 35, with a median of 26 days. Safety, eligibility, and symptom severity were assessed prior to enrolment by a registered consultant psychiatrist (co-author B.B.). All participants met criteria for treatment-resistant depression, defined as failure to respond to two or more adequate trials of pharmacological treatment and, where appropriate, psychotherapy.

### TMS protocol

Structural and resting-state functional MRI data (3T, 15 minutes eyes-open) were acquired prior to treatment to personalise stimulation targets. Participants received approximately 20–30 weekday sessions of intermittent theta burst stimulation^2,30,32^ (600 pulses at 120% resting motor threshold) while at rest, with treatment duration adjusted based on clinical response and practical considerations. Details can be found in Table 1 and the Supplementary Information.

### Symptom data acquisition

Depressive symptom severity before and after TMS treatment was assessed by a clinician using the Montgomery–Åsberg Depression Rating Scale (MADRS)^52^, while the severity of anxiety-related symptoms was assessed using the Hamilton Anxiety Rating Scale (HAM-A)^53^. In addition, the Hospital Anxiety and Depression Scale (HADS)^31^ self-report questionnaire was administered weekly throughout treatment. In the present analyses, we focused on the HADS depression subscale (HADS-D) because it provided repeated self-reported symptom measurements throughout the treatment course. As expected, scores in the clinician-administered scales were significantly correlated with self-report questionnaires (Supplementary Figure 1c).

### Medication and comorbidity

Information on participants’ medication and comorbidities is provided in the Supplementary Information.

### EEG data collection

An initial EEG protocol consisted of 7 minutes of resting-state EEG with alternating eyes-open and eyes-closed periods (2 minutes eyes open, 4 minutes eyes closed, 1-minute eyes open). Fifty participants completed this protocol. For the remaining 20 participants, the protocol was modified to include only 7 minutes of eyes-closed resting-state EEG (see suppl. fig. 11 for control analysis including EEG recording length). The analyses presented here focused exclusively on eyes-closed data. Accordingly, eyes-closed segments were extracted from the mixed-protocol recordings for the first 50 individuals, and the first and last minute of data were discarded for participants who completed the eyes-closed–only protocol to account for differences in recording length. EEG recordings were acquired immediately before and after TMS sessions 1, 11, and 20, resulting in six EEG recordings per participant (**Figure 1**). These sessions correspond to the beginning, midpoint, and the final EEG session. EEG data were recorded using a 64-channel ANT Neuro system with a Waveguard net electrode cap at a sampling rate of 1000 Hz.

### EEG preprocessing & source reconstruction

Raw resting-state EEG data were preprocessed using a fully automated pipeline implemented in a custom Python environment (version 3.12). The pipeline followed the recommended workflow of the Oxford Centre for Human Brain Activity and was implemented using the open-source *osl-ephys* (version 2.2dev0) package^33,34^, which builds on functionality from MNE-Python^35^ and FSL^54^. Data were band-pass filtered between 0.5 and 125 Hz and downsampled to 250 Hz. Bad channels and segments were identified and removed using the generalized extreme studentized deviate (G-ESD) algorithm^55^. Independent component analysis (ICA) was then applied to remove ocular artefacts. Subsequently, bad channels were interpolated, and data were re-referenced to the common average. In addition, all datasets were visually inspected prior to source reconstruction to confirm adequate data quality. Source reconstruction was performed using individual structural MRI scans and a linearly constrained minimum variance (LCMV) beamformer. Source-level data were parcellated using the Giles-38 cortical parcellation provided in *osl-ephys*, which covers the entire cortex. Spatial leakage was corrected using symmetric multivariate leakage correction^56^, and source polarity ambiguity was resolved via sign flipping based on lagged partial correlations^57^. Finally, parcel time courses were standardised.

### Canonical time-delay Hidden Markov Model, model inference, and summary statistics

We applied a canonical TDE-HMM to the parcellated time-series data for each session (10 states for the main analyses, 6 and 8 state model for control analyses), concatenated across subjects. The canonical model (v0.1+) was trained on the Cambridge Centre for Ageing and Neuroscience (Cam-CAN) dataset, which includes MEG resting-state and task data from 612 healthy participants aged 18– 88 years (1,849 recordings in total^58^). Details of model training are described in Gohil et al. (2026)^36^. See Supplementary Information for further details on the canonical TDE-HMM, model inference, and summary statistics.

### Transition probability PCA

State transition probability matrices were obtained for each participant and timepoint from the hidden Markov model. For each observation (participant × timepoint), we vectorized the transition probability matrix by extracting all off-diagonal elements (excluding self-transitions), yielding a feature vector of transition probabilities. Off-diagonal elements were used because self-transition probabilities were substantially larger and would otherwise dominate variance. Feature vectors were z-standardized across observations and subjected to principal component analysis (PCA). Participants/timepoints were then projected onto the PCA space to obtain component scores for the two components that explained the most variance (PC1/PC2). Component loadings were reshaped to the original state-by-state transition matrix format for visualization and interpretation. PCA was implemented in Python (version 3.12) using the *scikit-learn* package^59^.

### Statistical analysis

Baseline associations between brain state dynamics and depressive symptoms were assessed using ordinary least squares regression. FO at baseline (Session 1, pre-TMS) was analysed separately for each state, with values logit-transformed to account for the bounded range of the data. FO was regressed on baseline depressive symptom severity (HADS-D), controlling for age and gender, with heteroskedasticity-robust standard errors (HC3) and Benjamini–Hochberg false discovery rate (FDR) correction across states. To assess baseline transition dynamics, PCA component scores were regressed on baseline HADS-D, controlling for age and gender. To test whether stimulation-related changes predicted symptom improvement, pre–post changes in fractional occupancy (ΔFO = pre − post) were computed for each session and used to predict subsequent symptom severity (Session 1→11 and Session 11→20), controlling for symptom severity at the preceding timepoint, age and gender. The same regression framework was applied to PC2 and cycle metrics (cycle strength and rate). Regression coefficients are reported with standard errors and 95% confidence intervals, with significance set at α = 0.05 (two-tailed). Analyses were implemented in Python within the statsmodels package^60^.

### Temporal network density analysis (TINDA)

Temporal network density analysis (TINDA) was applied to characterise directed interactions between dynamic brain states^25^. Details regarding the implementation of the method are presented in the Supplementary Information.

## Code availability

Analysis code is available on [GitHub](https://github.com/CarinaFo/tms_brainpattern) in a public repository.

## Data availability

De-identified clinical data will be made available upon publication, following a DTA signed with QIMR Berghofer. EEG and fMRI data were collected under a commercial funding agreement and are not publicly available.

## Acknowledgments

We thank Oliva Heppell, Simon T. Thwaites, Jessica Miller, and Simon Issa, the QNC technicians, for providing us with high-quality EEG data. The authors thank P.B. for help with figures. This study was supported by the Australian NHMRC and MRFF (grant NHMRC2027597 to LC and NHMRC2021292 and MRFF2023308).

## Author contributions

C.F.: Conceptualization, Formal analysis, Software, Visualization, Writing – original draft. B.B.: Conceptualization, Data curation, Methodology, Resources. I.K.: Investigation. M.V.E.: Formal analysis, Interpretation of results. M.W.: Interpretation of results, Writing – review & editing. C.G.: Formal analysis, Investigation, Writing – review & editing., D.V.: Investigation, Writing – review & editing. M.V.H.: Writing review & editing. C.H.: Conceptualization, Formal analysis, Investigation, Funding acquisition, Writing – review & editing. L.C.: Conceptualization, Funding acquisition, Investigation, Supervision, Writing – original draft.

