## Supplementary Material for "Anterior default mode brain state dynamics predict depressive symptom severity before and during TMS treatment"

Supplementary Text

p. 1-2

Supplementary Figures

SFigure 1, p.3
SFigure 2, p.4
SFigure 3, p.5
SFigure 4, p.6
SFigure 5, p.6
SFigure 6, p.7
SFigure 7, p.7
SFigure 8, p.8
SFigure 9, p.9
SFigure 10, p.10
SFigure 11, p.11

Supplementary Tables

STable 1, p.12
STable 2, p.13
STable 3, p.14

Supplementary References

p.15

### Supplementary Text

**Symptom improvement**

Depressive symptom scores (HADS-D) were analysed using a linear mixed-effects model, with session treated as a categorical fixed effect and participant included as a random intercept (sessions: Session 1, Session 11, Session 20; supplementary analysis: pre, TMS Sessions 1, 11, 20, post).

**Medication and comorbidity data processing**

We extracted information on current and previous medications from clinical records. We identified individual free-text medication entries using regular expressions in Python. Next, each medication was mapped to a pharmacological class (STable 2). We report the number and percentage of participants taking at least one medication from each pharmacological class. The same process was adopted to map comorbid diagnoses, focusing on the most prevalent conditions (comorbidities present in more than one individual).

**TMS protocol**

Structural and resting-state functional MRI data (3T, 15 minutes, eyes-open) were acquired to personalise TMS therapeutic targets. The individual left dorsolateral prefrontal cortex (DLPFC) targets were defined based on fMRI signal anticorrelation with the subgenual cingulate cortex (SGC) using a connectivity-based approach[1]. TMS was delivered to the personalised brain target using neuronavigation (Localite) and a robotic arm (Axilum Robotics Cobot). The stimulation was delivered via a MagPro R30 machine and an actively cooled figure-of-eight coil (Cool-B65). The stimulation protocol for each TMS session was intermittent theta burst stimulation[2,3] (iTBS, 600 pulses at 120% resting motor threshold) while participants were at rest. Details regarding the whole neurostimulation procedure can be found in Burgher et al[4]. The treatment duration was adjusted based on clinical response and practical considerations, but the current study considered only TMS sessions during which EEG and clinical data were recorded (Sessions 1-20).

**Sample characteristics**

The sample was clinically heterogeneous. Thus, we also examined participants' clinical status by employing a validated grouping approach[5]. Research tier 1 included 15 individuals who would qualify for a randomised controlled trial (RCT) based on the consensus definition of treatment-resistant depression, whereby treatment resistance cannot be in part explained by the presence of complex psychiatric or neurological comorbidity or better explained by another primary disorder. Tier 2 included individuals (N = 45) with complex comorbidities who would not meet the RCT criteria. Research tier 3 included 10 individuals who did not meet the RCT criteria due to a structural neurological disorder or a diagnosis of bipolar depression. The most common comorbidities across the three tiers were attention-deficit/hyperactivity disorder and post-traumatic stress disorder (STable 1). Most individuals were taking antidepressants during TMS treatment, and only four participants were unmedicated (STable 2).

**Canonical time-delayed Hidden Markov Model**

We applied a canonical time-delay embedded HMM (TDE-HMM) to the concatenated time series for each individual across all sessions considered. The canonical model was trained on the Cambridge Centre for Ageing and Neuroscience (Cam-CAN) dataset, which includes MEG resting-state and task data from 612 healthy participants aged 18–88 years (1,849 recordings in total[6]). Details of model training are described in Gohil et al. (2025)[21]. The models are publicly available at https://github.com/OHBA-analysis/Canonical-HMM-Networks. Briefly, an HMM is a generative model that assumes a time series is generated by transitions among a finite set of mutually exclusive, recurrent, and temporary hidden brain states. In the TDE-HMM, each brain state is characterised by a unique spatio-spectral profile defined by patterns of signal power and coherence across brain regions.

Each time point is assigned to a single state, with the Markov assumption that the state at time *t* depends only on the state at time *t–1*.

**Model inference and spectral analysis**

We performed inference on our clinical EEG dataset by applying a canonical TDE-HMM pre-trained on the Cam-CAN data[7]. EEG data preparation followed Gohil et al. (2026)[7], including time-delay embedding (15 lags) and dimensionality reduction using principal components and covariance obtained from the canonical model. We used the same cortical parcellation and the same sampling frequency (250Hz) as the canonical model. The pre-trained model was applied to the concatenated dataset to infer posterior state probabilities (state time courses) across participants and sessions.
Next, we estimated state-specific spectral properties using multitaper spectral analysis applied to the non-embedded parcellated source EEG data. This was done using osl-dynamics functions[8] (default settings). Power spectra and coherence were estimated by weighting each time point by its posterior state probability, yielding state-specific measures. Spatial distributions of power and coherence were mapped onto cortical parcels to obtain state-specific spatio-spectral profiles.

**Brain-states time-courses**

After obtaining state-specific time courses for each individual and session, we calculated standard HMM summary statistics. These were:

- Fractional occupancy or FO: proportion of total time spent in each state
- Mean lifetime (ms): average duration of continuous state activation
- Mean interval (s): average duration between successive activations of a state
- Switching rate (Hz): average number of distinct state activations per second

Across summary measures, brain states exhibited distinct profiles, with no single state dominating the state space (SFig. 3). Each state had a unique topology and spectral profile (see SFig. 2 for visualisation of the states' EEG signal power topology and cross-parcel coherence).

**Brain state matching for confirmatory analyses**

We compared state-specific coherence profiles between the 10-state solution and alternative 6- and 8-state solutions. For each session, group-level coherence matrices were computed by averaging subject-level coherence estimates using the subject-specific spectral weights. Coherence was then averaged and vectorised using the upper triangle of the parcel-by-parcel coherence matrix. We then computed Pearson correlations between the vectorised coherence profiles from the original states and each state profile from the alternative HMM solutions. The state with the highest correlation was taken as the best-matching state for that model resolution and session. Overall, fractional occupancy, transition probability, and cycles results replicate across resolutions (e.g., SFigure 8, 9, 10).

**Temporal network density analysis (TINDA)**

Temporal network density analysis (TINDA) was applied to characterise directed interactions between dynamic brain states[9]. For each state $m$, inter-state intervals were divided into first and second halves, and a $K \times K$ fractional occupancy asymmetry matrix was computed. Each matrix element quantified the difference in occupancy of state $n$ between the two halves of state $m$'s inter-state intervals. This matrix was interpreted as a weighted, directed graph capturing systematic directional biases in state transitions and was computed separately for each participant and session. Cycle strength was quantified to assess whether state transitions formed a coherent cyclical structure. To estimate cycle rate, a time-resolved measure of state visit counts was derived using a sliding window equal to the mean state lifetime. A second-level HMM with Poisson observations, constrained to follow the identified cycle, was then fitted. Cycle duration was defined as the time required to complete one full cycle, and its inverse (cycle rate) was used in statistical analyses. All TINDA analyses were performed using the Python implementation provided in *osl-dynamics (version 2.1.dev8)*[8] and custom functions.

#### Supplementary Figures

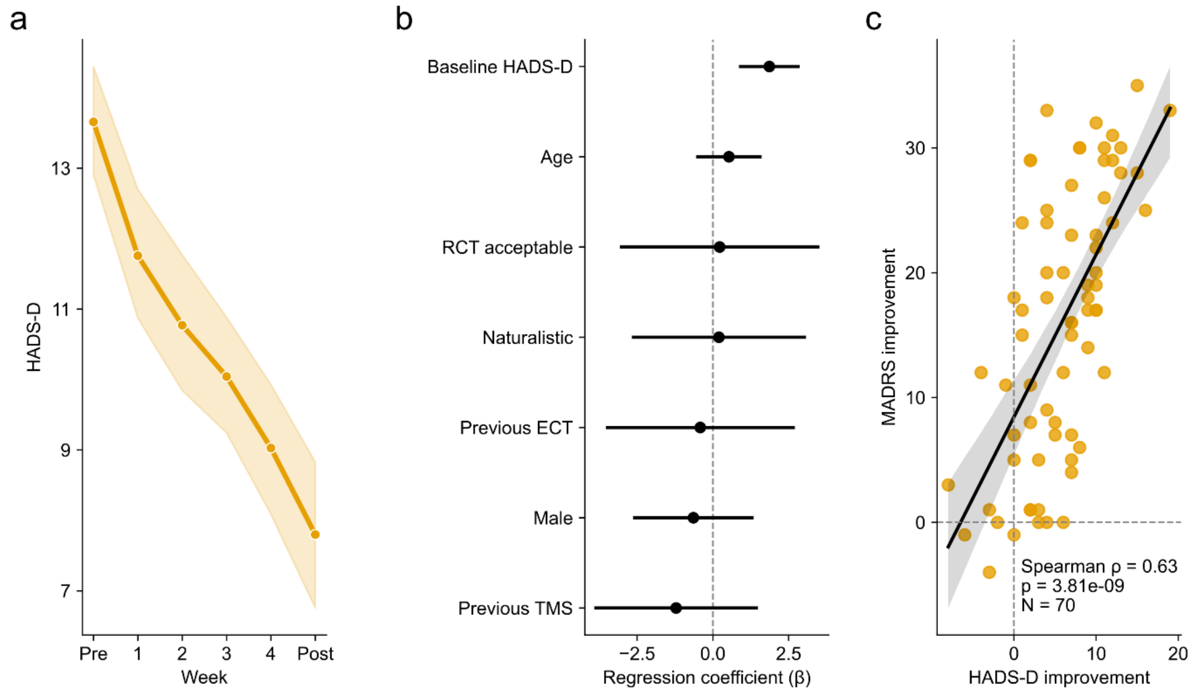


**SFigure 1.** *Symptom change during TMS treatment, baseline predictors of symptom improvement, and correlation between clinician-administered and self-reported depressive symptoms*. **a**. Symptom trajectory over the course of TMS treatment shows an improvement in the severity of depressive symptoms from pre to post treatment ($p < 0.001$ for each session; Hospital Anxiety and Depression scale-Depression, HADS-D[10]). The plot shows the mean HADS-D score per timepoint, and the shaded area represents the standard error of the mean (N=70). **b.** Baseline participant characteristics did not significantly predict the HADS-D score at Session 20 (all p>0.2). **c.** Spearman correlation between symptom changes (baseline *minus* post assessments depression score ) measured using the HADS-D (self-report) and the Montgomery–Åsberg Depression Rating Scale (MADRS[11], clinician administered, N=70).

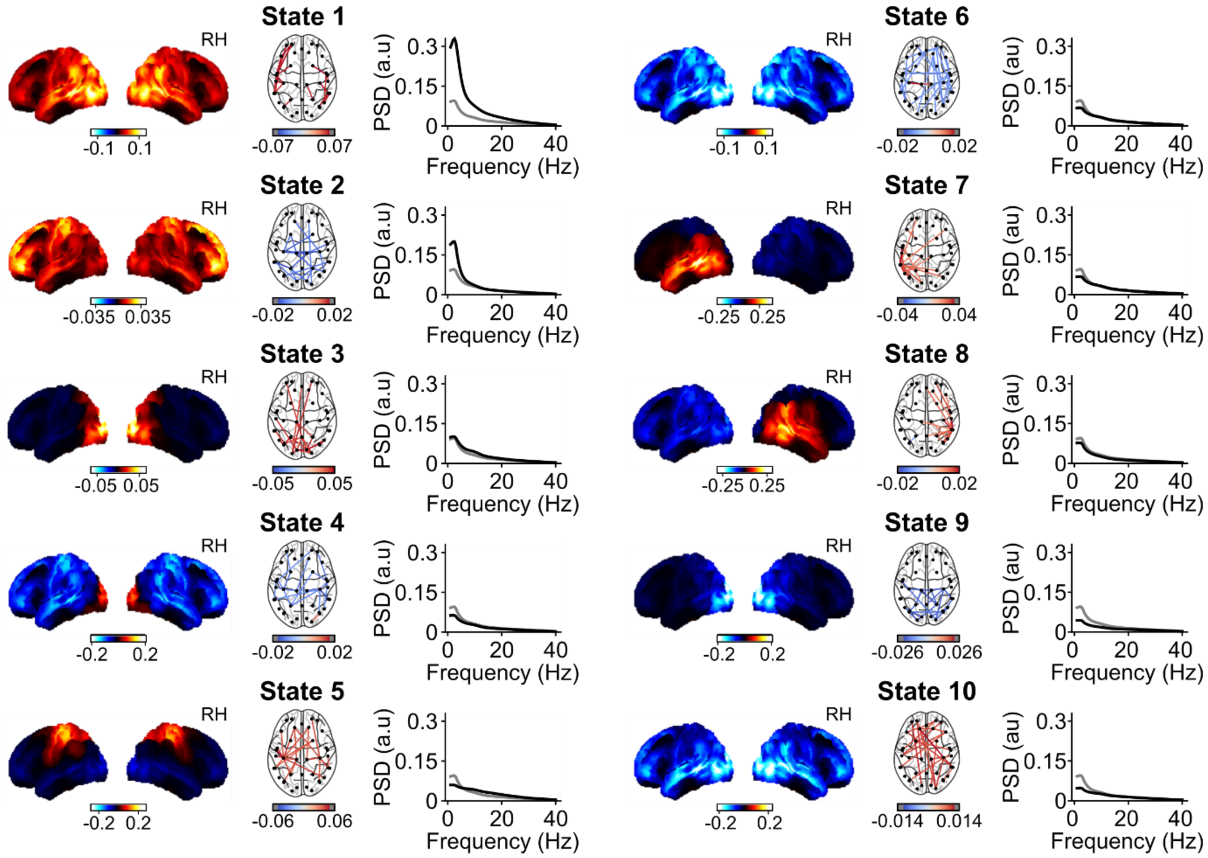


**SFigure 2.** *Brain states defined by the 10-state canonical TDE-HMM model*. Eyes closed resting-state EEG data collected during TMS treatment (6 sessions) for depression were modelled using a 10-brain-state canonical time-delay embedded Hidden Markov Model (TDE-HMM)[12]. The model was pre-trained on the CAM-CAN dataset[6]. Each brain state is represented by a relative power map (1-40Hz), a coherence network (1-40Hz, showing the top 3 % of connections), and a power spectral density (PSD) profile. The figure shows the average output over 70 individuals at baseline (i.e., first EEG recording before any TMS). States 3 and 4 are dominated by occipital activity and are referred to as visual brain states. State 5 shows the highest power in the somatosensory cortices, whereas State 6 shows relatively low power and coherence in and between the visual, parietal, and temporal regions. Brain states 7 and 8 are centred on the left and right temporal lobes and commonly referred to as auditory states, while states 9 and 10 exhibit reduced posterior power and have been associated with attentional processing[13,14].

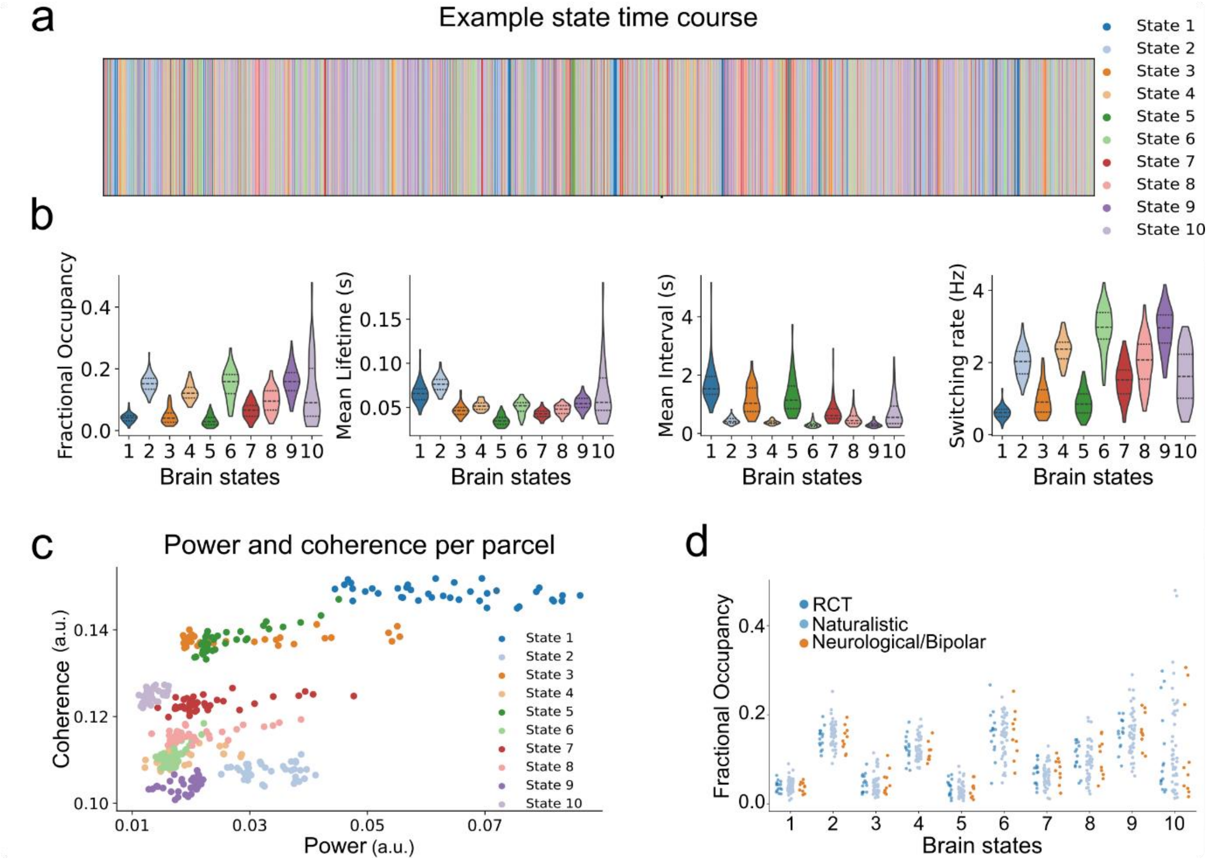


**SFigure 3.** *TDE-HMM summary statistics*. **a.** Representative brain-states time course (10-state model fitting). The x-axis represents time, with only one state active at any given time. **b.** HMM summary statistics of resting-state brain state dynamics before any TMS (baseline- i.e., Session 1 pre, see Figure 1a in the main text): Violin plots for fractional occupancy (FO), mean lifetime, mean interval rate and switch rate for each brain state. **c.** Mean EEG signal power and coherence per cortical parcel for each state. **d.** Research groups according to consensus classification framework[5]: RCT (tier 1), participants eligible for randomized clinical trials; Naturalistic (Tier 2), participants with complex psychiatric comorbidities; Neurological (Tier 3), participants with a diagnosis of bipolar depression or a neurological condition. The panel depicts fractional occupancy for each brain state and participant at baseline, as a function of research group assignment. There are no differences in FO across the three groups.

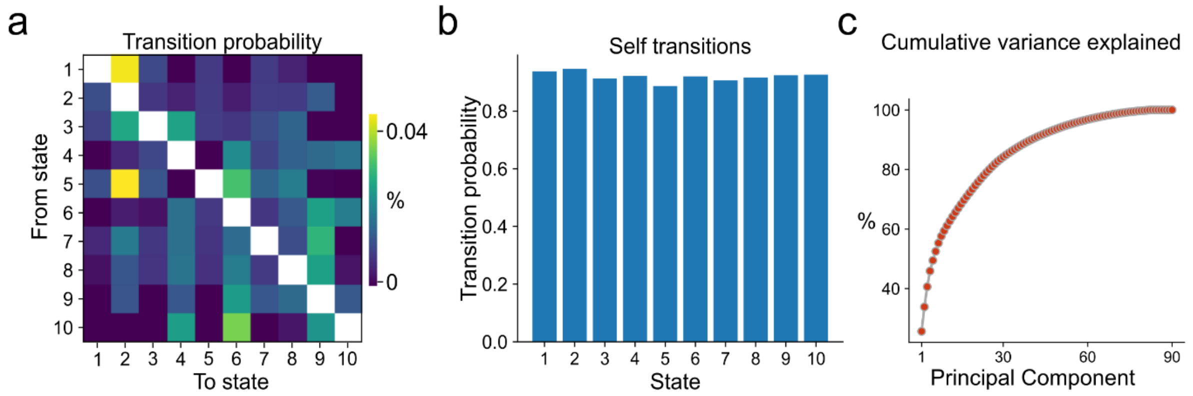


**SFigure 4.** *Brain state transition probabilities.* **a.** Transition probability matrix for the baseline EEG session (Session 1, pre-TMS; Figure 1a in the main text) averaged over participants, excluding self-transitions (diagonal). **b.** Self-transitions obtained from the transition probability matrix averaged over participants at baseline. **c.** Cumulative variance explained by a Principal Component Analysis (PCA) performed on the transition probability (excluding self-transitions) matrices of each participant and session (input array dimensions: N_sessions, N_participants, N_states, N_states).

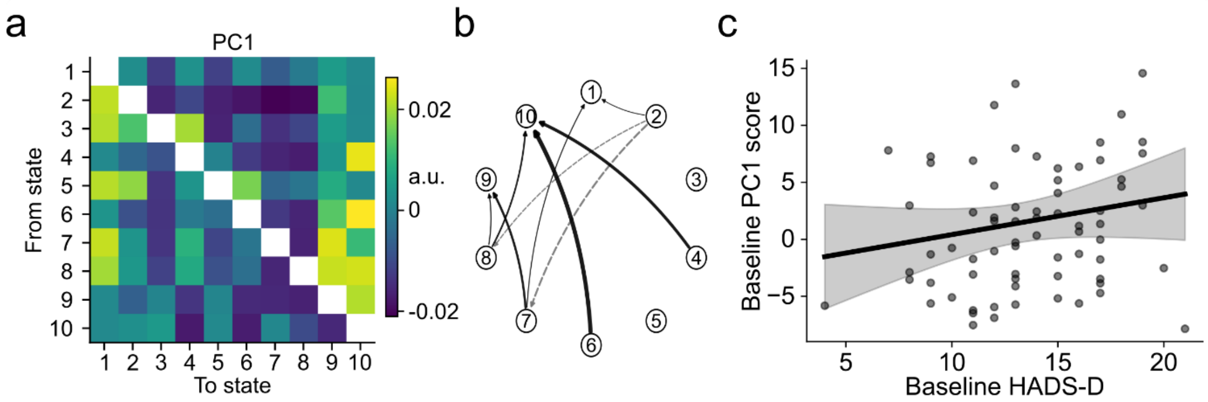


**SFigure 5.** *PC1 on the transition probability matrix does not significantly predict baseline depression severity***.** The first principal component (PC1) on the baseline (Session 1 pre-TMS) transition probability matrices shows strong positive loadings on transitions into attention-related brain states (states 9 and 10, SFigure 2) and negative loadings on transitions out of the anterior default mode state (state 2). **b.** Graphical presentation of the 20 % absolute strongest transitions. **c.** Baseline depressive symptom severity (HADS-D) did not significantly predict the PC1 score in the first EEG session ($p_{uncorrected} = 0.173$).

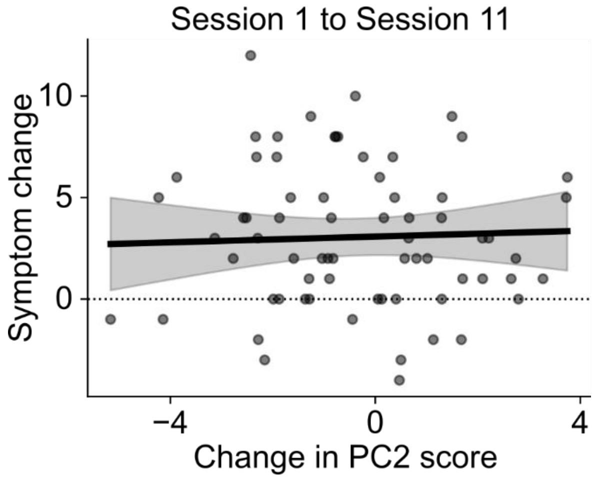


**SFigure 6.** *The change in PC2 in Session 1 does not significantly predict symptom change from Session 1 to Session 11***.** The change in PC2 score from pre to post TMS in Session 1 (Figure 1a) did not significantly predict symptom change in self-reported HADS-D scores from Session 1 to Session 11 ($p_{uncorrected} = 0.803$).

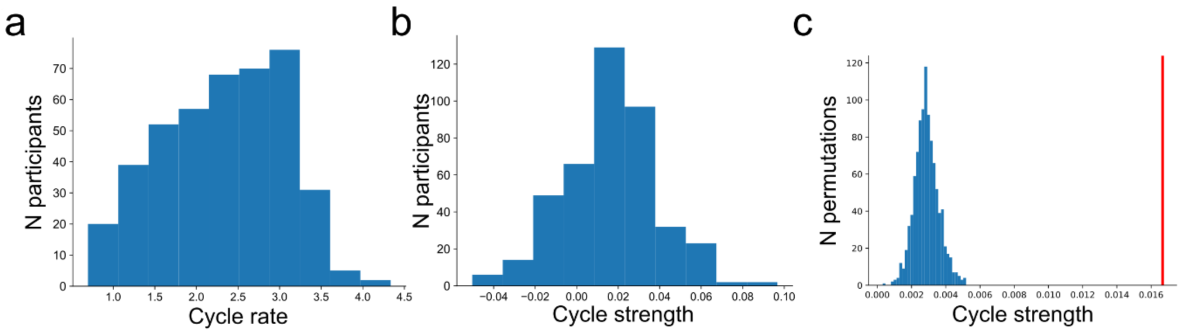


**SFigure 7.** *Cycle features.* **a.** Histogram of cycle rate (Hz) over the six EEG sessions (Figure 1a) and participants. **b.** Histogram of cycle strength (a.u.) over all sessions and participants. **c.** Permutation statistic on observed *versus* permuted cycle strength. To assess statistical significance, we generated a null distribution by randomly permuting the state labels within each participant and session 1000 times, while preserving the temporal structure of the state time courses. For each permutation, cycle strength was recomputed. The observed cycle strength was then compared to the distribution of cycle strengths obtained from the permuted data. The observed cycle strength is highlighted by the red vertical line ($p < 0.001$).

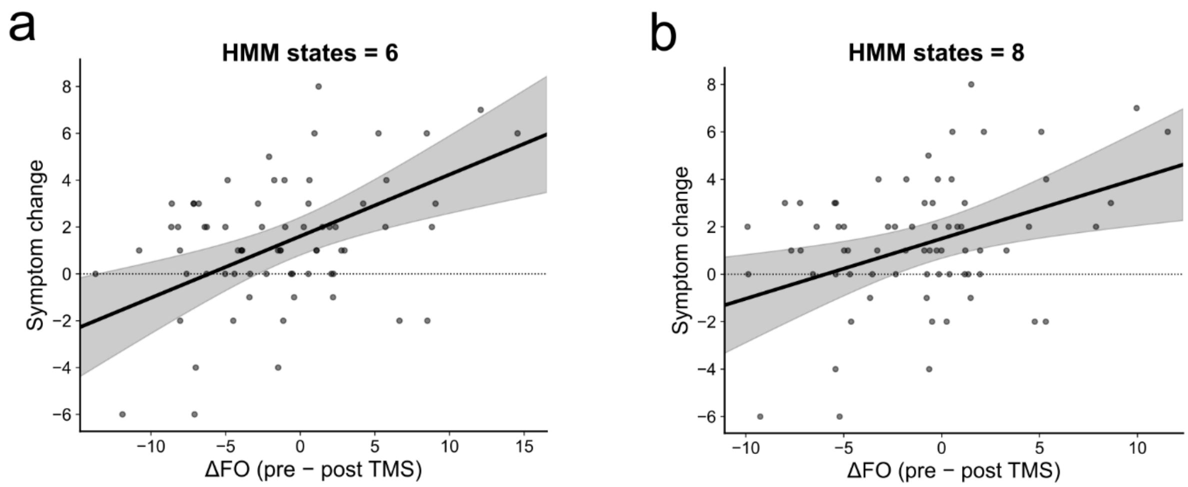


**SFigure 8.** *Control analysis: Brain state resolution on fractional occupancy.* Validation of the predictive value of changes in the anterior default mode state pre *versus* post TMS (delta FO) for symptom changes using a 6-state and 8-state TDE-HMM canonical model. We matched state 2 in the 10-state model to the state with the highest correlation based on spatial coherence (state 2 in all model resolutions; Pearson $r > 0.98$). The relationship between delta FO and symptom improvement at mid-treatment is significant for an HMM resolution of fewer than 10 states. Changes in anterior default mode FO (states 2 and 10) significantly predict symptom improvement from session 11 to session 20 in a 6-state HMM (a, anterior default mode state = state 2, $p < 0.001$) and a 8-state HMM (b, anterior default mode state = state 2, $p = 0.004$).

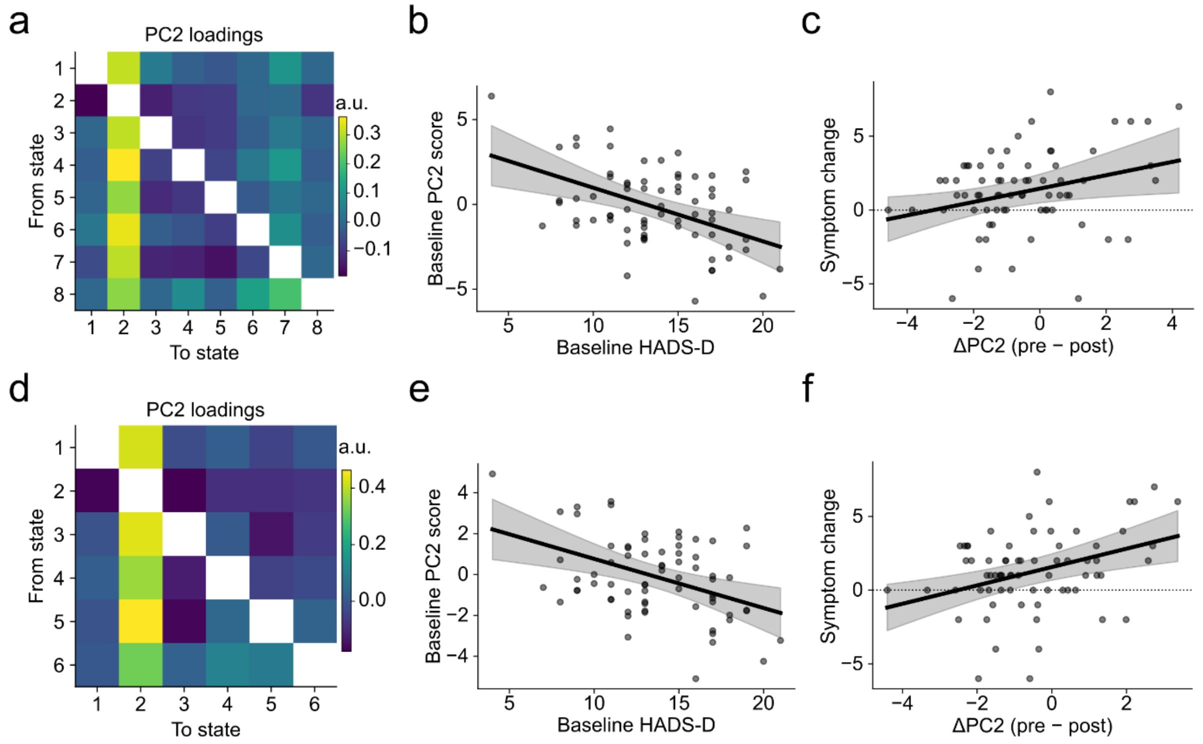


**SFigure 9.** *Control analysis: Brain state resolution on transition probabilities.* Validation of the predictive value of changes in transition probabilities pre *versus* post TMS (delta PC2) for symptom changes using a 6-state and 8-state TDE-HMM canonical model. We matched state 2 in the 10-state model to the state with the highest correlation based on spatial coherence (state 2 in all model resolutions; Pearson r > 0.98). a. The second principal component has strong positive loadings on transitions into state 2 (anterior default mode state) in the 6 (a) and (d) 8 states HMM. b. A higher baseline HADS-D score predicts a lower PC2 score in Session 1 in the 8 (c, p<0.001) and 6 states (e, p=0.001) HMM. c. The relationship between delta PC2 (pre minus post TMS) and symptom change from Session 11 to Session 20 is significant in the HMM with 8 states (c, p=0.017) and 6 states (f, p=0.001).

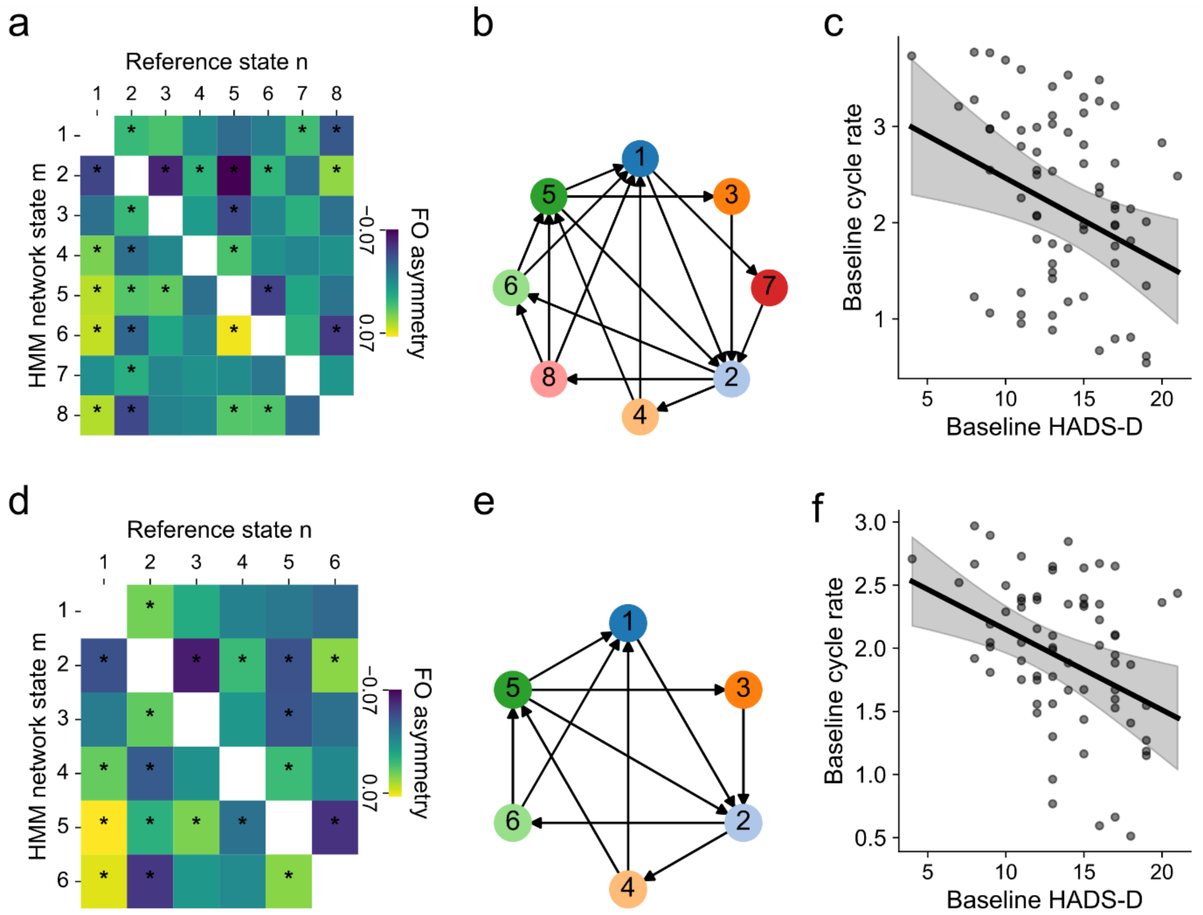


**SFigure 10.** *Control analysis: Brain state resolution on cycle rate.* Validation of the predictive value of differences in cycle rate and baseline depressive symptom severity using a 6-state and 8-state TDE-HMM canonical model. a. The fractional occupancy asymmetry matrix shows significant asymmetries (highlighted with an asterisk, Bonferroni corrected at a=0.05) for the 8 (a) and (d) 6 states HMM. b. Optimised cyclical order for the 8 states HMM (b) and the 6 states HMM (e). c. Baseline HADS-D significantly predicts cycle rate in Session 1, with a higher cycle rate for participants with lower HADS-D scores in the 8 states HMM (c, p=0.022) and the 6 states HMM (f, p=0.003).

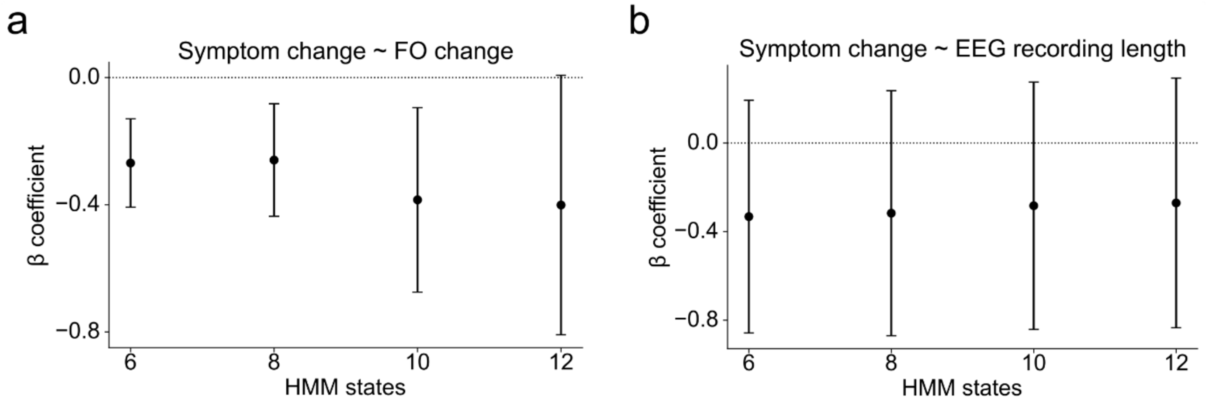


**SFigure 11.** *Control analysis: EEG recording length.* To ensure that variability in EEG recording duration (min=162s, max=320s, mean=216s) did not bias the results, EEG recording length was included as a covariate in all regression models predicting symptom change from Session 11 to Session 20 based on pre-post TMS change in fractional occupancy obtained from State 2 (anterior default mode state). EEG recording length was averaged across pre- and post-TMS recordings within Session 11 and standardised (z-scored). The inclusion of this covariate did not alter the pattern or significance of the main effect of FO change (a, $p_{6states} < .001$, $p_{8states} = 0.004$, $p_{10states} = 0.009$, $p_{12states} = 0.054$), and the EEG recording length was not a significant predictor of symptom change (panel b; all $p > 0.2$).

#### Supplementary Tables

**STable 1.** *Psychiatric comorbidities.* The table summarises the number and percentage of participants diagnosed with a comorbid psychiatric disorder. It includes only diagnoses reported by more than one participant. Five participants had no comorbid disorder.

| **Psychiatric Comorbidities** | **n (%)** |
|---|---|
| Attention-deficit/hyperactivity disorder | 28 (43.1%) |
| Post-traumatic stress disorder | 20 (30.8%) |
| Generalised anxiety disorder | 12 (18.5%) |
| Autism spectrum disorder | 9 (13.8%) |
| Eating disorder | 7 (10.8%) |
| Obsessive-compulsive disorder | 7 (10.8%) |
| Alcohol use disorder | 4 (6.2%) |
| Social anxiety disorder | 2 (3.1%) |
| Tourette syndrome | 2 (3.1%) |

**STable 2.** *Previous and current medications.* The table summarises the number and percentage of participants taking at least one medication in each class (both current and previous). Information on one individual's current medication and on the previous medications of three individuals was missing.

| **Medication class** | **Current medications n (%)** | **Previous medications n (%)** |
|---|---|---|
| Antidepressants | 50 (71.4%) | 64 (91.4%) |
| ADHD agents | 25 (35.7%) | 2 (2.9%) |
| Antipsychotics | 13 (18.6%) | 16 (22.9%) |
| Benzodiazepines | 11 (15.7%) | - |
| Mood Stabilisers | 10 (14.3%) | 5 (7.1%) |
| Unmedicated | 4 (5.7%) | - |

**STable 3.** *Symptom changes over the course of TMS treatment.* Clinician-administered MADRS and HAM-A scores were collected at baseline and at Session 20. Self-reported HADS scores were collected weekly during treatment.

| **Timepoint** | **MADRS** | **HAM-A** | **HADS-D** | **HADS-A** |
|---|---|---|---|---|
| **Baseline** | 36.0 (33.0–40.0) N=70 | 27.0 (19.0–35.0) N=69 | 13.0 (11.0–16.8) N=70 | 15.0 (12.0–17.0) N=70 |
| **Week 1** | – | – | 11.0 (9.0–14.8) N=70 | 13.0 (10.0–15.0) N=70 |
| **Week 2** | – | – | 10.0 (8.0–13.0) N=70 | 12.0 (9.2–15.0) N=70 |
| **Week 3** | – | – | 9.5 (8.0–12.0) N=70 | 11.0 (9.0–13.8) N=70 |
| **Week 4** | – | – | 8.0 (7.0–11.0) N=70 | 10.0 (8.0–13.8) N=70 |
| **Post** | 18.0 (10.0–27.5) N=70 | 15.0 (7.0–24.0) N=69 | 8.0 (4.2–10.0) N=70 | 10.0 (7.0–13.8) N=70 |
| **Change Baseline → Post** | -17.0 (-25.0–-7.0) N=70 | -8.0 (-17.0–-2.0) N=69 | -6.5 (-10.0–-2.0) N=70 | -3.5 (-7.0–-1.0) N=70 |

Values represent the median over participants and the interquartile range.
N indicates the number of participants with available data at each time point.
